# Gut microbiota metabolically mediate intestinal helminth infection in Zebrafish

**DOI:** 10.1101/2024.07.26.605207

**Authors:** Austin J. Hammer, Chris A. Gaulke, Manuel Garcia-Jaramillo, Connor Leong, Jeffrey Morre, Michael J. Sieler, Jan F. Stevens, Yuan Jiang, Claudia S. Maier, Michael L. Kent, Thomas J. Sharpton

## Abstract

Intestinal helminth parasite (IHP) infection induces alterations in the composition of microbial communities across vertebrates, although how gut microbiota may facilitate or hinder parasite infection remains poorly defined. In this work we utilized a zebrafish model to investigate the relationship between gut microbiota, gut metabolites, and IHP infection. We found that extreme disparity in zebrafish parasite infection burden is linked to the composition of the gut microbiome, and that changes in the gut microbiome are associated with variation in a class of endogenously-produced signaling compounds, N-acylethanolamines, that are known to be involved in parasite infection. Using a statistical mediation analysis, we uncovered a set of gut microbes whose relative abundance explains the association between gut metabolites and infection outcomes. Experimental investigation of one of the compounds in this analysis reveals salicylaldehyde, which is putatively produced by the gut microbe *Pelomonas*, as a potent anthelmintic with activity against *Pseudocapillaria tomentosa* egg hatching, both *in vitro* and *in vivo*. Collectively, our findings underscore the importance of the gut microbiome as a mediating agent in parasitic infection and highlights specific gut metabolites as tools for the advancement of novel therapeutic interventions against IHP infection.

**Importance:** Intestinal helminth parasites (IHPs) impact human health globally and interfere with animal health and agricultural productivity. While anthelmintics are critical to controlling parasite infections, their efficacy is increasingly compromised by drug resistance. Recent investigations suggest the gut microbiome might mediate helminth infection dynamics. So, identifying how gut microbes interact with parasites could yield new therapeutic targets for infection prevention and management. We conducted a study using a zebrafish model of parasitic infection to identify routes by which gut microbes might impact helminth infection outcomes. Our research linked the gut microbiome to both parasite infection, and to metabolites in the gut to understand how microbes could alter parasite infection. We identified a metabolite in the gut, salicylaldehyde, that is putatively produced by a gut microbe and that inhibits parasitic egg growth. Our results also point to a class of compounds, N-acyl-ethanolamines, which are affected by changes in the gut microbiome and are linked to parasite infection. Collectively, our results indicate the gut microbiome may be a source of novel anthelmintics which can be harnessed to control IHPs.

## Introduction

Intestinal helminth parasitic infections present a significant global health burden, affecting at least one quarter of the global population^1,2^, and are disproportionately experienced by individuals in impoverished nations, particularly children^3^. These infections are also prominent among domestic animals^4,5^, which places tremendous strain on livestock management and veterinary practices^4,6,7^. Among infected individuals intestinal helminth parasite (IHP) infections can contribute to anemia^8^, cognitive impairment^9^, physical wasting^10^, as well as a host of other conditions that contribute to the equivalent of millions of disability-adjusted life years^11^. Unfortunately, the extreme burden of IHP infection may be exacerbated by the emergence of drug-resistant parasites. High levels of broad anthelmintic drug resistance in animal populations have been observed globally for decades^12,13^ posing a tremendous risk to humans because practices such as broad blanket anthelmintic administration^14^ and prior widespread prophylactic administration of anthelmintic drugs^15^ have provided strong selective forces for virulent drug resistant organisms^16^. While improved helminth management practice may help slow the rate of anthelmintic resistance^16,17^, the future of controlling IHP infection may depend on innovating new methods and resources in anthelmintic discovery to stay ahead in the pugilistic battle with drug-resistance in infectious nematodes.

In the search for new approaches to control intestinal helminth parasites (IHPs), there is suggestive evidence that the intestinal microbiome can enhance or reduce parasite infection^18^. Microorganisms produce a diverse trove of metabolic compounds, including anthelmintic drugs. For example, the avermectin class of compounds, which include the most widely administered anthelmintic drugs on the planet, were originally derived from soil-borne bacteria such as *Streptomyces avermectilis*^19^. Besides the soil, locations where helminths and microbes have evolved to co-locate, such as the gut, may offer a rich resource of microbially derived anthelmintic compounds^18^ as their evolution by microbial community members may have been critical to microbial exploitation of the shared ecological niche. However, the complex and variegated metabolic landscape of microbially derived compounds which are found in the gut and relevant to IHP infection remains unexplored.

Little is known about the existence of anthelmintic compounds derived from gut bacteria, but it is known that gut microbiota can drive intestinal helminth infection through a variety of mechanisms. For instance, bacteria can alter the protective integrity of mucosal barriers^20–23^, drive peristaltic activity^24,25^, and produce inhibitory antibiotic compounds that limit pathogen and parasite survival^19,26–28^. Moreover, signals from specific gut bacteria can cause egg hatching of helminthic parasites in the gut, such as in the case of *Trichuris muris* infections in mice ^29^. Additionally, gut microbes engage in extensive interaction with the vertebrate host immune system^30^, and intestinal helminths possess a diverse suite of immunomodulatory tools^31,32^. Much of this cross-talk depends on the production of metabolite products by host, microbe, and parasite. However, our current understanding of the complex set of metabolite interactions that may directly or indirectly drive parasite colonization success is limited. Clarifying the set of metabolites that mediate the interaction between host, parasites, and the gut microbiome may provide a toolkit of compounds to control parasite infection.

Efforts to understand the role of the gut microbiome in health and disease conditions can benefit from the application of analytic techniques that examine possible mediating roles of intestinal microbes and metabolites. To this end, mediation inference techniques seek to examine whether the relationship between two variables depends on the hypothesized mediating effect of some third variable. While early iterations of these methods relied on regression-based structural equation modeling^33^ with strict assumptions regarding model type, new methods are being innovated that account for data-specific assumptions, such as sparsity and compositionality in microbiome data^34–37^, and the suite of mediation tools available to researchers is expanding rapidly. Recent applications of mediation analysis in microbiome science have identified lipid compounds produced by *Akkermansia muciniphila* that modulate murine immunity and metabolism^38^, clarified the role of gut microbiota in mediating the relationship between diet and immune inflammation^39^, and established the role of the gut microbiome in the development of childhood asthma^40^. However, the extension of mediation techniques to high dimensional multi-omic data where there are a large number of both independent and mediating features remains limited. Large sample sizes are often required to attain sufficient power to detect mediating effects^41^, which can impose substantial logistical challenges on researchers seeking to use these techniques with expensive vertebrate models. Despite this challenge, mediation techniques have played an important recent role in mapping the gut microbiome to metabolites which may be involved in disease conditions such as anorexia nervosa^42^, and these methods offer a dynamic opportunity to ascertain the intricate mechanisms by which the gut microbiome links to complex diseases, such as parasite infection.

Robustly identifying novel connections between the gut microbiome and parasite infection via mediating metabolites presents a challenge that requires experimental investigation using an organism that displays robust patterns of quantifiable IHP infection, displays a tractable set of gut microbes, and may be scaled in the lab to deal with inherent statistical limitations of mediation methods. In line with these requirements, the zebrafish model provides a powerful tool for modeling parasite infections and shedding light on the intricate relationship between host-microbiota interactions^43^ and disease outcomes^44,45^. The model possesses a well characterized taxonomic gut microbial composition^46,47^ with a functional composition which resembles that of humans^48^, zebrafish offers a well-established model of intestinal helminth parasite infection^49^, and can be experimentally scaled in a cost-effective manner^50–52^. Zebrafish have previously been used to discover and assay disease-related natural products^53–55^ with broad relevance. Collectively, the zebrafish-IHP model can be highly controlled to investigate intricate routes by which the gut microbiome may mediate parasite infection, and insights gleaned from connections between gut microbiota and gut metabolites may offer translational potential for understanding the metabolite-based interactions between the gut microbiota, intestinal helminth parasites, and the host.

In order to uncover gut microbial metabolites that mediate IHP infections, we used a zebrafish model of intestinal helminth infection by the nematode *Pseudocapillaria tomentosa*^49,56^, as it affords access to the large sample sizes needed to disentangle these relationships. In particular, instead of housing fish together we individually-housed parasite infected fish hosts to understand why a small number of zebrafish bear a disproportionate parasite burden in the absence of social or co-housing dynamics, and produced paired microbiome and metabolome data from infected and uninfected fish to investigate links between the microbiome, parasitic infection, and intestinal metabolites. We observed that the gut microbiome explains the variation in infection burden across individuals. We then utilized a mediation inference framework to identify microbe-metabolite interactions that statistically mediate worm burden. This work reveals a potent anthelmintic, salicylaldehyde, whose effect on infection burden is mediated by members of the gut microbiome. Analysis of the paired microbiome-metabolome data also implicates N-acylethanolamines (NAEs) in the association between microbiota and parasite infection burden. Collectively, our work demonstrates that the zebrafish gut microbiome metabolically mediates IHP infection outcomes and reveals novel microbiome-sourced anthelmintic drug leads.

## Results

### Intestinal helminthic parasite infections are overdispersed among socially isolated zebrafish

IHP infections frequently manifest overdispersed distributions across wildlife populations, agricultural settings, and scientific laboratories^57–59^. Prior work has shown that social behavior and interactions can drive differences in parasite infection burden^60–62^ but it remains unclear if such behavior and interactions underlie the distribution of burden. For example, in zebrafish, social hierarchies and behaviors^63–65^ may impact feeding, which could result in interindividual biases in oral exposure to infectious agents. To understand if zebrafish parasite burden overdispersion occurs in the absence of co-housing or social dynamics, we individually housed 100 zebrafish in 1.2-L tanks and exposed 50 fish to *P. tomentosa* eggs. Stool samples were collected from all surviving individuals at several time points: prior to *P. tomentosa* exposure, immediately before exposure, and 29 days following parasite exposure, which prior work has shown is the peak of infection^49,66^. At this final time, point fish were sacrificed and infection burden was quantified through cytological analysis of dissected intestinal tissue.

Interrogation of the distribution of infection burden among *P. tomentosa*-exposed fish 29 days post exposure (dpe) reveals that infection is overdispersed across the population (σ²/μ=4.719, Supplementary Figure 1), indicating that relatively small numbers of exposed individuals carry the bulk of mature worms in their guts. Our investigation of socially isolated individuals reveals that community social dynamics alone are not the sole drivers of overdispersed helminth parasite worm burden among zebrafish populations and indicates that other factors underlie this phenomenon.

### Gut microbiome composition associates with parasite exposure and infection burden

Gut microbiomes display highly personalized forms across individuals^67,68^, and variation in gut microbial communities has been consistently linked to parasite infection^66,69,70^. Thus, it is conceivable that overdispersion in parasite infection burden outcomes results from the intricate interplay between parasite exposure and gut microbiome composition that occurs in each individual, where bacterial consortia could tip the scales toward susceptibility or resilience. We investigated if zebrafish gut microbiome community structure is related to parasite exposure and subsequent overdispersion of parasite infection burden. To investigate how gut microbiome composition changes before and after parasite exposure, we generated 16S rRNA gene sequence data from stool samples collected at both a pre-exposure baseline and at 29 days post-exposure (dpe). In order to test whether the initial microbial community state influences the microbiome’s association with infection burden at the peak of infection, we initially altered fish gut microbial communities by administering antibiotics three days prior to *P. tomentosa* parasite exposure (Fig. 1). This strategy was employed to ensure variability in the gut microbiome compositions across the individually housed fish, which was necessary for analyzing the potential role of the gut microbiome in parasite infection. The resultant microbial diversity enabled us to explore the association between distinct microbiome profiles and the differential parasite burden outcomes. Microbial communities at 29 dpe were significantly stratified by both parasite exposure (PERMANOVA, F=7.1618, p<0.0001) as well as parasite burden (PERMANOVA, F=12.1514, p<0.0001). These results are consistent with our earlier findings^66^, but in this case individually housing fish eliminates possible impacts of co-housing that may drive homogeneity of the gut microbiome among infected versus non-infected individuals. Thus, this design and these results provide particularly compelling evidence that zebrafish gut microbial communities are connected to *P. tomentosa* and later infection success, and may contribute to the overdispersion of infection burden across individuals.

**Figure 1.**
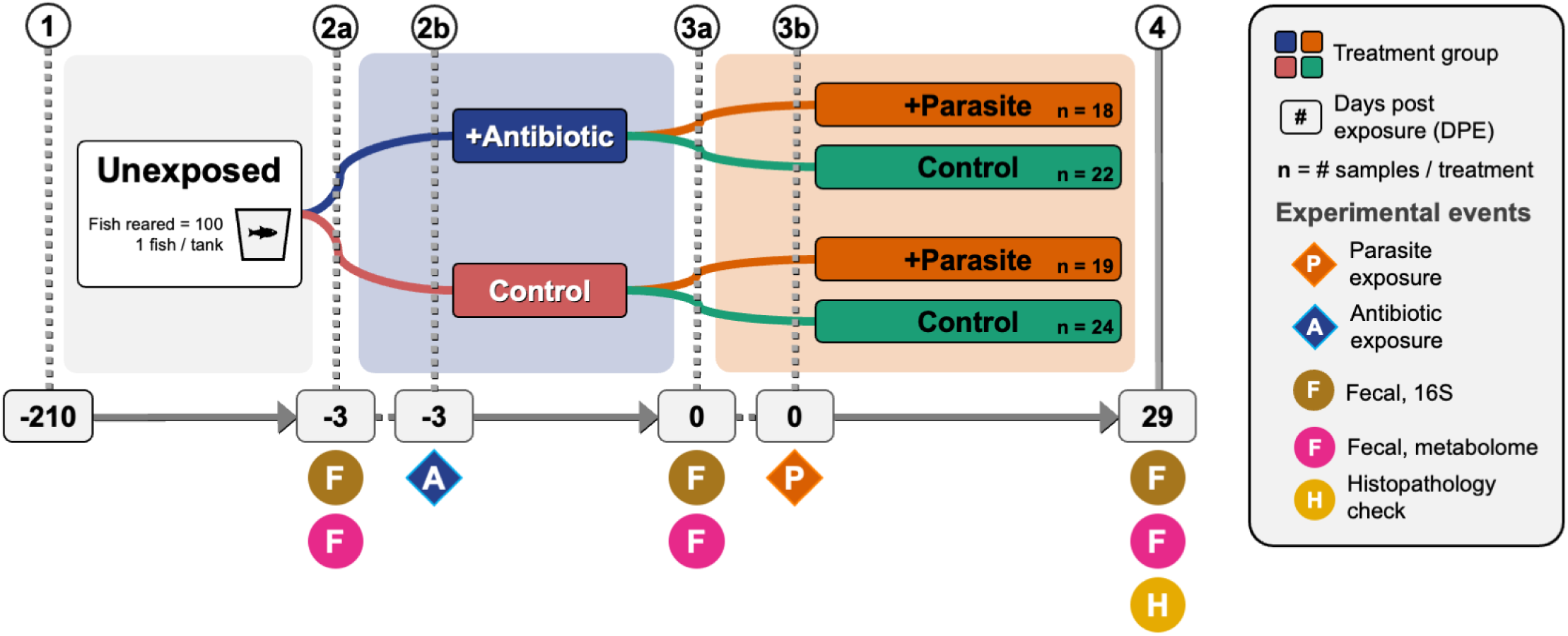
Schematic of zebrafish husbandry and treatment events and timeline. 1) Briefly, 100 adult fish were placed in individual tanks, 2b) half of fish were subsequently exposed to antibiotics, 3b) then fish were randomly exposed to the zebrafish parasite Pseudocapillaria tomentosa. Fecal samples were collected 2a) prior to antibiotic exposure, 3a) just prior to parasite exposure, and 4) 29 days post-parasite exposure (dpe) after which fish intestinal histopathology was assessed. Samples were split and processed for untargeted fecal metabolomic analysis as well as fecal 16S rRNA DNA amplicon sequencing.

### Effect of IHP exposure on the gut microbiome composition depends on the pre-exposure microbiome state

Given that parasite colonization and the effects of parasite infection have been linked with gut microbiome composition in this work and elsewhere^71^, we reasoned that altering the initial state of the microbiome may shape its relationship to subsequent IHP infection. Analysis of 16S rRNA gene sequences generated from fish stool collected after this antibiotic exposure but prior to *P. tomentosa* exposure reveals that antibiotic administration successfully altered the composition of the zebrafish gut microbiome (PERMANOVA, F=27.565, p<0.0001). A corresponding analysis at 29 dpe shows that the relationship between fecal microbial community composition and IHP exposure depends on this prior antibiotic exposure (PERMANOVA, F=3.16, p=0.009). Moreover, we find that parasite infection burden is strongly linked to the composition of the microbiome (PERMANOVA, F=7.1618, p<0.0001, Fig. 2), but that this relationship fundamentally depends on whether hosts were exposed to antibiotics first (PERMANOVA, F=4.2087, p=0.002). This interaction is particularly noteworthy because at 29 dpe no strong relationship is observed between the composition of the gut microbiome and antibiotic exposure alone (PERMANOVA, F=1.32, p=0.2277). These findings collectively suggest that perturbations to the initial state of the microbiome, such as through antibiotic exposure, have a cryptic effect on the successional interplay between IHP infection and the gut microbiome, even when the statistical effects of antibiotic exposure are no longer prominently observed.

**Figure 2.**
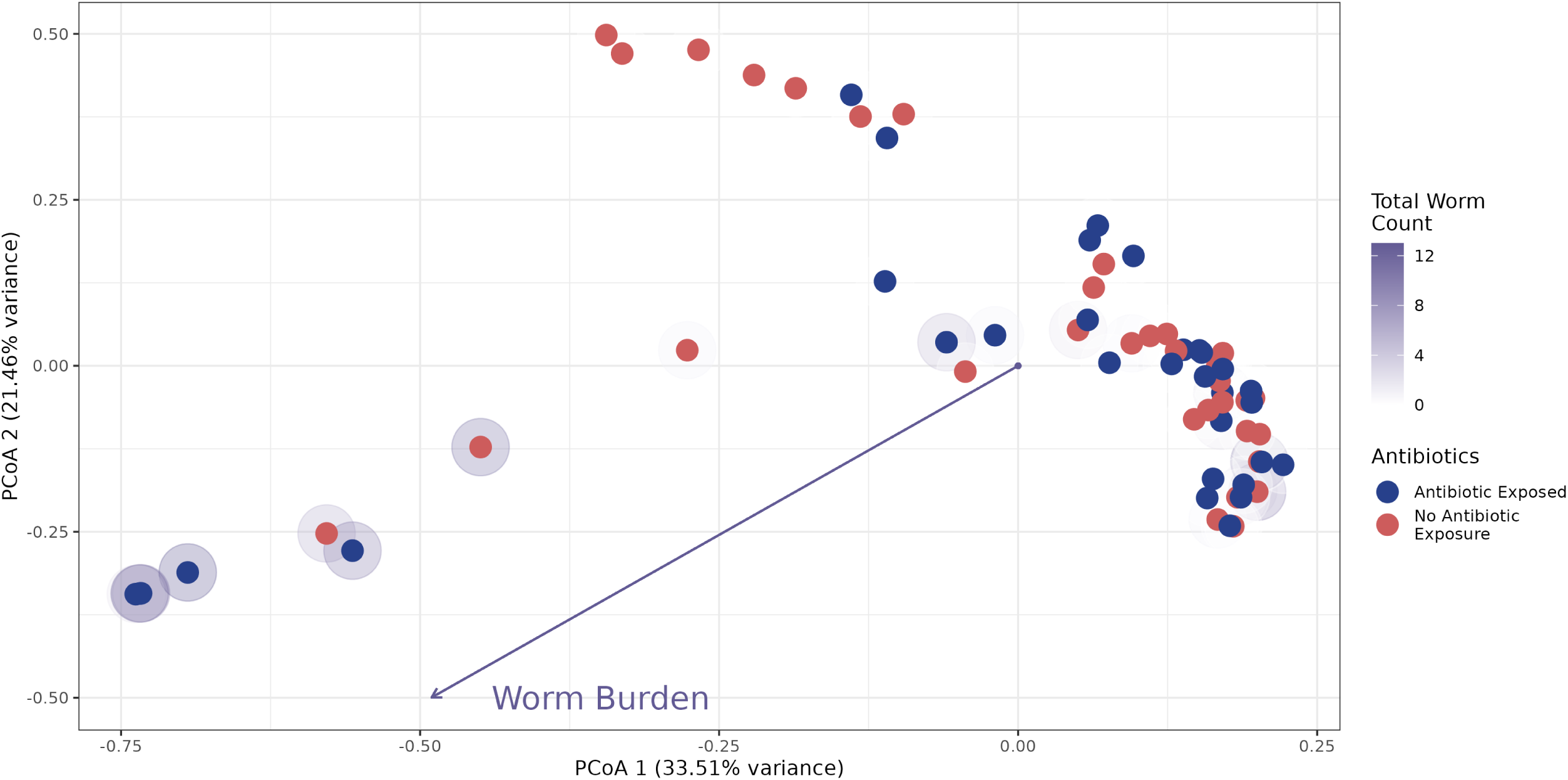
Principal Coordinates Analysis (PCoA) ordination of Bray-Curtis dissimilarity of microbiome communities at 29 days following parasite exposure. Each point represents an individual fish. The halo intensity around points represents the number of quantified parasites in the gut at dissection. Point colors represent antibiotic exposed (blue) and unexposed (red) groups. The arrow illustrates an envfit relationship for worm burden, depicting the linear direction of association between parasite burden and Bray-Curtis dissimilarity.

This observation may be of special interest given that geographic locales which have higher levels of IHP infection also tend to be locations where microbiomes may be disrupted by the use of antibiotics to manage bacterial infections^72,73^. Given the varied influential roles of the microbiome on parasite infection^26–29^, disruption of the initial microbiome state by antibiotics may interfere with the ability of the microbiome to protect the host from helminth infection. Elucidating how antibiotic use alters the microbes involved in helminth resistance, as well as their specific interactions with parasites, is crucial. This knowledge could guide the development of microbiome-based therapies that supplement those protective elements and reduce the adverse impacts of helminth infection.

### Fecal metabolites, including salicylaldehyde and N-acylethanolamines, inversely associate with helminth infection burden

Metabolic products convey information about the presence of pathogens and microbes^70,74^, produce signals that underlie immune control^75^, and interact within a complex network of gut microbes, the host immune system, and invading parasites, driving shifts in the intestinal environment. In order to understand the metabolomic landscape wherein zebrafish gut microbes and parasites co-locate, we performed untargeted metabolomic profiling of the fecal samples collected from the same individuals and at the same time point as our microbiome profiling analysis. Prior research has shown that the zebrafish gut metabolome is composed of a diverse array of lipids and fatty acids^76,77^, as well as amino acids and various biogenic amines^78,79^. Consistent with previous research, our annotated gut metabolomic data is also dominated by a large number of complex lipids, vitamin and amino acid derivatives, as well as polar metabolites from many compound classes. We first sought to find metabolites from this diverse metabolite set that are statistically associated with parasitic worm burden. Due to the high level of overdispersion frequently observed in parasite infection data^57–59^ we utilized negative-binomial generalized linear models (GLMs) to examine the statistical relationship between a set of 303 annotated metabolites and *P. tomentosa* worm burden. We uncovered 35 metabolites which associate with parasite worm burden among infected hosts (Negative Binomial GLM, FDR<0.1, Supplementary Table 1).

Numerous compounds which are associated with helminth parasite worm burden have also been linked to parasite infection in other work. Salicylaldehyde, a compound previously noted as a soil and plant nematicide^80–82^ shares an inverse relationship with worm burden. This analysis also identifies compounds which have previously been linked to parasite infection such as a major form of Vitamin E, gamma-tocopherol^83^. Additionally, at 29 dpe we find that 6 of the 8 compounds classed as N-acylethanolamines (NAEs) such as oleoyl ethanolamide (OEA), linoleoyl ethanolamide (LEA), 2-linoleoyl glycerol, and related N-acylethanolamine (NAE) precursor compounds (e.g., glycerophospho-N-oleoyl ethanolamine), manifest an abundance profile that sharply distinguishes infected versus uninfected individuals, where infected individuals display higher metabolite abundances (Wilcoxon-Rank Sum Test, p<0.05, Fig. 3a). Furthermore, six of the eight NAE-related compounds in these data are strongly inversely associated with parasite worm burden (Negative Binomial GLM, FDR<0.1, Fig. 3b, Supplementary Table 1).

**Figure 3.**
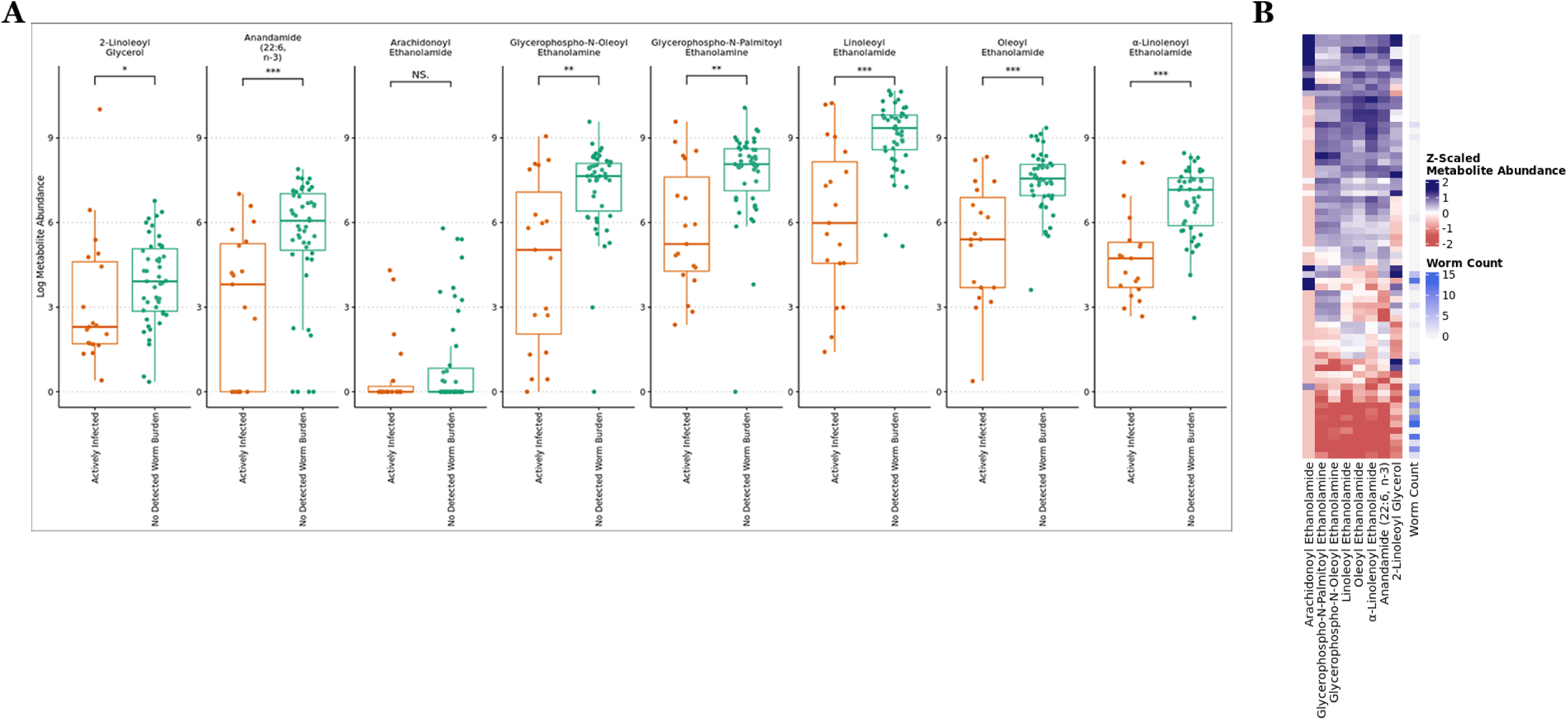
(A) The abundance of N-Acyl-Ethanolamine related (NAE) compounds significantly differs between infected and uninfected fish. (B) NAE abundance inversely associates with parasite exposure for six of eight identified compounds. “*” indicates p<0.05, “**” p<0.01, and “***” p<0.001.

The NAE class of metabolites represents a broad family of lipid messengers that play a well-established role in energy metabolism and feeding behavior^84–86^, as well as inflammation^85^, and prior work has established a relationship between the gut microbiome and NAEs^87–89^, so we next tested if the abundance of these metabolites is related to microbiome composition and antibiotic exposure. Strikingly, we observed a robust association between the abundance of these NAE compounds and gut microbiome composition at a time point prior to parasite exposure and following antibiotic treatment. (PERMANOVA, p<0.05, Supplementary Table 2). Notably, at this time there is evidence that the relationship between the zebrafish gut microbiome and the abundance of six NAEs depends on whether or not fish were exposed to prior antibiotics (PERMANOVA, p<0.1, Supplementary Table 2). These results are further underscored by our finding that the composition of the gut microbiome at 29dpe is still strongly linked to NAE abundance for six different compounds (PERMANOVA, p<0.0002, Supplementary Table 2). Furthermore, the relationship between the gut microbiome and NAEs is underscored by our results that show antibiotic exposure interacts with NAE abundance in a manner that is significantly related to gut microbiome composition for five of the eight NAE metabolites (PERMANOVA, p<0.0002, Supplementary Table 2).

These findings reinforce the emerging view that the gut microbiome plays a fundamental role in regulating NAE levels, an observation which is especially noteworthy because recent work has shown that intestinal nematodes which infect humans, mice, and even insects have genes that encode functions for degradation of NAEs^90^. The collective impact of our NAE analysis shows that their abundance is starkly different in parasite-uninfected versus infected hosts, that NAE abundance is linearly related to parasite infection burden, and that the gut microbiome is a principal driver of NAE abundance in zebrafish hosts. Given the multifaceted role of NAEs in host physiology, immune response, and intestinal microbiome control, plus the ability of IHPs to degrade and produce these compounds, emphasizes the significance of these compounds as a nexus in the battle between helminth parasites and vertebrates host.

### Connections between fecal metabolite abundance and *P. tomentosa* worm burden are putatively mediated by fecal microbiota

The prior analysis linking metabolites to parasite worm burden highlighted several compounds which are also known to drive changes in intestinal microbiome composition. Given the complex interplay between gut microbes and metabolite production, these findings open a line of inquiry into microbiome-metabolite interactions. Zebrafish gut microbial taxa have been linked to parasite infection^66^, and given the connection between some of the aforementioned metabolites, such as NAEs, we hypothesized that the relationship between members of the gut microbial community and parasite infection depends on metabolite-related cross talk. In order to explore the interconnected role of metabolites, microbiota, and parasite worm burden we selected parasite burden-linked metabolites and prevalent taxa, then statistically analyzed individual metabolite-microbe pairings to identify relationships which may be relevant to parasite burden. Our workflow applied partial correlation to a set of worm burden linked metabolites and prevalent ASVs in order to identify microbe-metabolite associations which were robust after accounting for the controlling influence of other microbial taxa. Additionally, we used mediation inference methods to quantify whether a metabolite’s statistical relationship to worm burden may be hypothetically mediated by members of the microbiota. The results of this approach provided a set of potential interactions between 25 metabolites and 17 members of the microbiota (Adjusted Causal Mediating Effect FDR<0.3, Fig. 4), whose microbe-metabolite relationship is uniquely strong in the context of other ASVs, and whose interacting relationship may be relevant to parasite infection.

**Figure 4.**
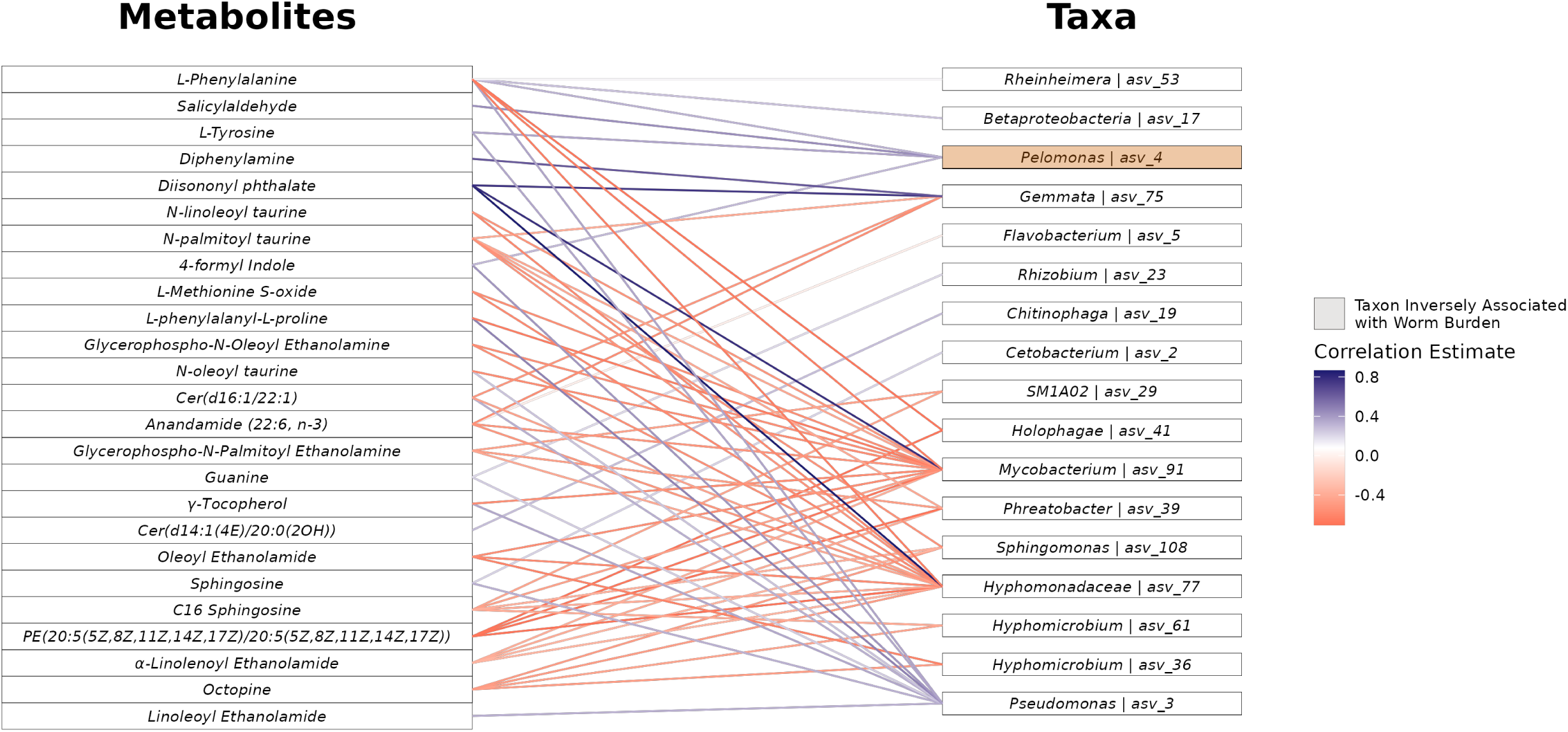
Network of microbe-metabolite interactions predicted to mediate parasite worm burden. Nodes represent fecal metabolites and gut bacteria. Edges represent statistically significant relationships, with colors indicating the direction of correlation (blue: positive, red: negative).

One particular edge in this possible interaction set includes salicylaldehyde and an amplicon sequence variant (ASV) from the *Pelomonas* genera. Salicylaldehyde, also known as 2-hydroxybenzaldehyde, is an organic compound that occurs naturally in some foods such as buckwheat^91^ and it is known that some salicylaldehyde derivatives exhibit antibacterial activity^92^. As described earlier, salicylaldehyde manifests a robust inverse relationship among parasite exposed fish 29 days following parasite exposure (Fig. 5a) and the microbe to which it is correlated, ASV 4, is one member of a small set of microorganisms in this work that are negatively related to total helminth worm burden (Negative Binomial GLM p=0.002, Fig. 5b). In addition to this inverse relationship following parasite infection, we also find that it is a particularly important feature for predicting later parasite infection burden. We constructed a random forest regression model considering parasite worm burden as a function of all gut microbial taxa present prior to parasite exposure, and found that *Pelomonas*, specifically ASV 4, shows up as one of the most important taxa for predicting parasite worm burden among a feature set that includes hundreds of different microbial taxa (Supplementary Figure 2). This same ASV is also strongly correlated with salicylaldehyde abundance (Spearman’s Correlation, ρ=0.62, p=0.002, Fig. 5c) and is predicted to mediate the relationship between salicylaldehyde and worm burden (Average Causal Mediation Effect (ACME) FDR<0.3). Little is known about the function and metabolic capacity of *Pelomonas*, but given its relationship with salicylaldehyde and its strong inverse relationship with helminth worm burden we surveyed available *Pelomonas* genomes available in the NCBI genome repository to understand if the taxonomic group possesses genetic pathways associated with salicylaldehyde metabolism. Several available *Pelomonas* genomes possess salicylaldehyde dehydrogenase^93^, an enzyme responsible for metabolism of salicylaldehyde, typically as part of the naphthalene degradation pathway^94^, and it has been long demonstrated that some taxa from the *Pelomonas* genera are capable of metabolizing diverse polycyclic aromatic hydrocarbons (PAHs) including naphthalene^95^. *Pelomonas* metabolic capacity to interact with salicylaldehyde could very well explain the predicted mediating role of this taxon in the relationship between salicylaldehyde and worm burden.

**Figure 5.**
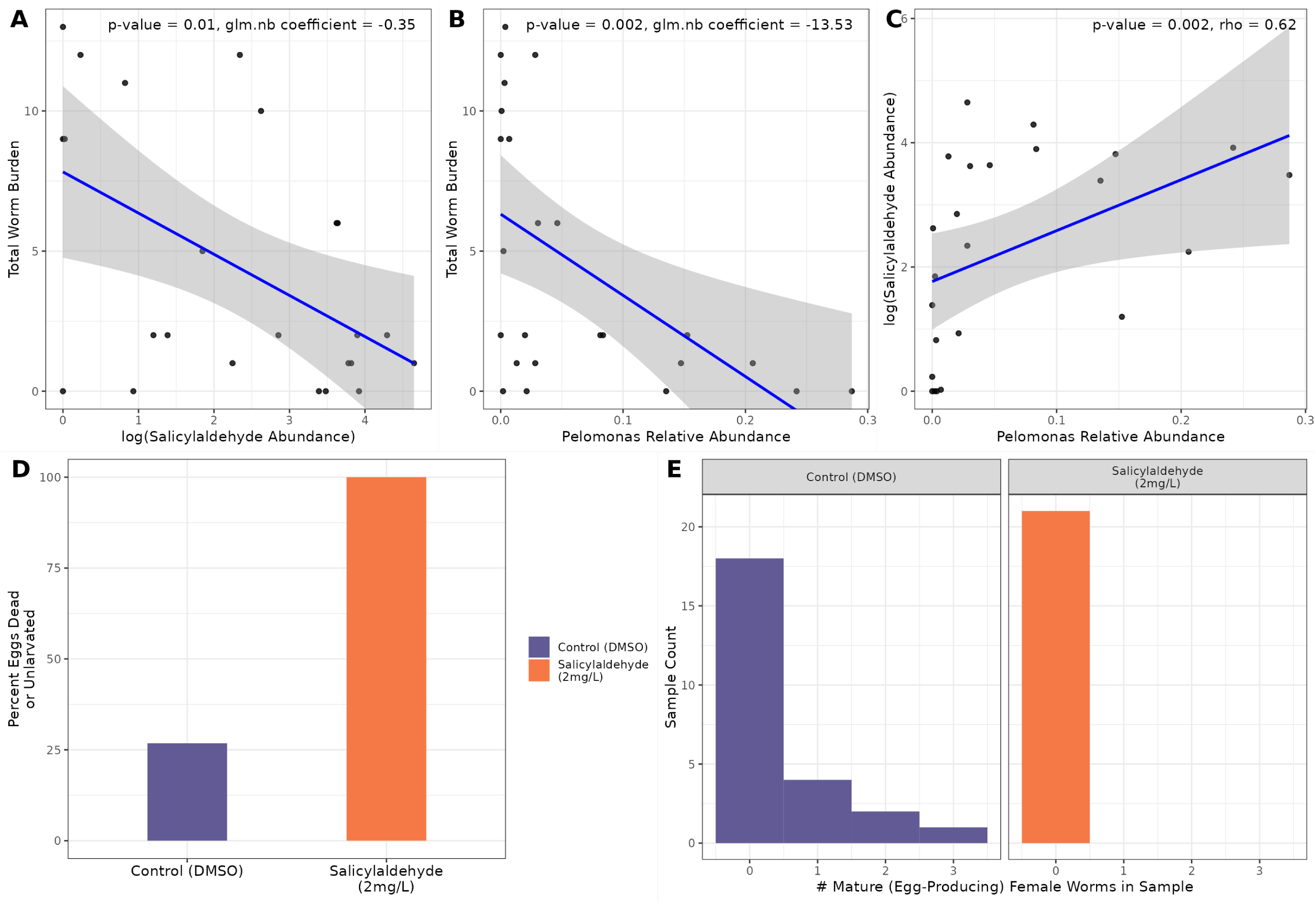
(A) Scatterplot of log salicylaldehyde abundance against worm burden among fish that are 29 dpe. (B) Scatterplot of *Pelomonas* ASV 4 relative abundance against worm burden among parasite exposed fish 29dpe. (C) Linear relationship between log salicylaldehyde abundance and *Pelomonas* relative abundance. (D) *P. tomentosa* eggs were exposed to salicylaldehyde at a concentration of 2mg/L in an *in vitro* assay. The y-axis depicts the % of eggs that are unlarvated or dead. (E) *in vivo* salicylaldehyde exposure assay comparing the number of mature female *P. tomentosa* worms that produced eggs.

We also found that the relationship between γ-tocopherol, which is a major form of vitamin E, and parasite worm burden is putatively mediated by ASVs from the *Mycobacterium* and *Pseudomonas* genera and parasite worm burden. The abundance of these ASVs was positively linked to worm burden in this work (Negative Binomial GLM, FDR<0.05), and in an earlier investigation the genera-level abundance of both of these taxa displayed a positive correlation with IHP burden in zebrafish^66^. γ-tocopherol has been shown to modulate the composition of the gut microbiome^83,96^ and mitigate colitis caused by LPS-induced inflammation signaling in mice^97^, and reduced abundance of this metabolite leaves mouse hosts more susceptible to helminth infections^98,99^. If this compound displays similar colitis mitigating and gut microbiome altering effects in zebrafish, reduced abundance of this metabolite, either by parasite infection or through metabolism by parasite-promoting taxa, may help clarify the basis through which these members of the microbiota associate with increased worm burden.

### Salicylaldehyde administration completely inhibits *P. tomentosa* egg larvation

To determine the anthelmintic effect of salicylaldehyde, we utilized both *in vitro* and *in vivo* drug exposure assays to determine how exposure to the drug impacts *P. tomentosa*. One mechanism common to many anthelmintic compounds such as albendazole and ivermectin is inhibition of worm egg larvation^100^. Therefore, we innovated an *in vitro* assay to evaluate how salicylaldehyde exposure impacts *P. tomentosa* egg larvation. Specifically, exposed 131 *P. tomentosa* eggs to salicylaldehyde at a dose of 2mg/L, 136 eggs at 15mg/L, and a group of 242 eggs that were reared without exposure to salicylaldehyde. For each group, we quantified larvation rates after 5 days. In the salicylaldehyde-unexposed group, 174/242 (72%) of the eggs successfully larvated. Conversely, 0 eggs from either salicylaldehyde treated groups larvated (Fisher’s Exact Test, p<2e-16, Fig. 5d). The complete inhibition of egg larvation demonstrates notable inhibition of egg-hatching in *P. tomentosa*.

We followed up on this finding by determining if salicylaldehyde can disrupt active *P. tomentosa* infection *in vivo*. To do so, we exposed 60 fish to *P. tomentosa* parasite eggs, then split the fish equally into 30 fish which were exposed to salicylaldehyde at a concentration of 2mg/L, and 30 fish that were not exposed to salicylaldehyde, but to a DMSO control. Mortality in salicylaldehyde exposed and unexposed host fish was 10 and 5 fish, respectively, which started at 16dpe. All examined moribund or fresh dead fish showed infections. The experiment was terminated at 24 dpe, and the intestines of all surviving fish showed 100% infection in both groups. Results indicate a slight reduction in infection burden among salicylaldehyde exposed fish, with an average of 15 worms/fish in salicylaldehyde treated fish versus an average of 19 worms/fish among untreated controls (Wilcoxon Rank-Sum Test, p=0.3; Fig. 5e). Although more strikingly, we found that drug treatment interfered with worm development: 7/25 (28%) the fish that were not exposed to salicylaldehyde contained gravid female worms with eggs in their guts, whereas no female parasites or free eggs were recovered from the salicylaldehyde-exposed treatment group (Fisher’s Exact Test, p=0.01, Fig. 5e).

Overall, these assays indicate that salicylaldehyde represents an enticing anthelmintic drug lead, although further work is required to clarify the mechanism by which it acts, as well as to explore specificity of IHP types it is active against. Furthermore, elucidating the potential salicylaldehyde-producing capability of *Pelomonas* or other microbes may offer a microbiome oriented route for control of parasite infection.

## Discussion

The rise of anthelmintic resistant IHPs presents an exigent challenge to identify new drugs and techniques to control IHP infection. To do so we must expand our understanding of the factors that underlie infection, as this knowledge can be leveraged to design new biocontrol strategies. Based on the historical discovery of novel anthelmintic compounds among bacteria^19^ as well as accumulating evidence that the intestinal microbiome affects the colonization success of IHPs^18,69^, we reasoned that exploring the metabolic landscape of the gastrointestinal microbiota in the context of infection could unlock new leads in the quest for novel anthelmintic strategies. To explore this idea, we investigated the relationship between gut microbes, metabolites, and IHP infection in a zebrafish model using a multi-omic approach to statistical mediation. In so doing, we identified a variety of metabolites that associate with infection burden in a manner that is dependent upon specific microbial taxa. These relationships are valuable to resolve not only because the metabolites and taxa in question may be utilized to develop novel infection control strategies (e.g., anthelmintic drugs or probiotics), but because they underscore the putative mechanisms by which the microbiome influences IHP infection outcomes.

Experimental validation of one particular lead, salicylaldehyde, reveals that our approach can uncover microbially mediated compounds that elicit anthelmintic effects. Salicylaldehyde is a nematicide used in agricultural settings and in our data is among the metabolites that are most strongly inversely associated with worm burden. We performed follow up *in vitro* and *in vivo* tests to confirm that salicylaldehyde elicits effects against *P. tomentosa*. In particular, this compound completely inhibits *in vitro P. tomentosa* egg development and maturation. Cessation of egg production following anthelmintic treatment is a common phenomenon with gastrointestinal nematodes in production animals. For example, moxidectin has been demonstrated to inhibit egg production in female worms of *Cooperia* that survive treatment in the early stages of resistance^101^. We observe a similar phenomenon here, where there is a complete absence of parasite worm eggs recovered from hosts that are exposed to salicylaldehyde. Previous work has also shown that salicylaldehyde prevented hatching of the potato cyst nematode, *Globodera pallida*^81^, but this compound is not currently used for the control of helminth infections in animal populations, and our experiments are the first to demonstrate that this compound is also capable of inhibiting egg maturation in a parasite which infects vertebrate organisms. While the precise mechanism of this action is uncertain, *P. tomentosa* eggs have relatively delicate shells and are quite susceptible to desiccation and chemical agents^102^. Thus, it is possible that salicylaldehyde either damages the egg shell directly, or possibly translocates to unlarvated worms where it may impair their larvation physiology. Future work should seek to uncover the specific mechanisms of action and whether salicylaldehyde can elicit broad effects across IHP species that infect other hosts. Regardless, our study suggests that repurposing salicylaldehyde as an anthelmintic drug against vertebrate IHPs may help control infection in the face of rising multi-drug resistant IHPs.

As zebrafish are not capable of producing salicylaldehyde, we used statistical mediation to identify a microbe from the *Pelomonas* genus as a potential source of the compound. In particular, our mediation analysis finds that the relative abundance of *Pelomonas* positively correlates with the abundance of salicylaldehyde and significantly explains the variation in the underlying relationship between the compound and infection burden. While a variety of processes may underlie these associations, prior work supports the hypothesis that these patterns result from *Pelomonas*-induced metabolism of salicylaldehyde. The *Pelomonas* genus is known to possess enzymes that may aid in the metabolism of salicylaldehyde, such as salicylaldehyde dehydrogenase^93^, and is a member of a diverse set of *Betaproteobacteria* which are known to generate and utilize this compound during the degradation of naphthalene^94^. While degradation of naphthalene represents one parsimonious explanation for the origin of salicylaldehyde, another important alternative hypothesis regarding the origin and synthesis of salicylaldehyde begins with phenylalanine. The results of our analysis point to phenylalanine as a metabolite which is inversely related to parasite worm burden and whose relationship with parasite worm burden is potentially mediated by *Pelomonas* (Fig. 4). The conversion of phenylalanine to trans-cinnamic acid is known to be performed by bacteria^103–105^. The following in a phenylalanine to salicylaldehyde metabolism might first require conversion of trans-cinnamic acid to o-coumaric acid. The terminal conversion of o-coumaric acid to salicylaldehyde is known in tobacco plants^106^, as well as in a species of fungus^107^, although bacterial catalysis of this terminal reaction is not characterized. While this might represent a parsimonious explanation for the biosynthesis of salicylaldehyde, more work is warranted to establish the distribution of microbial participation in these pathways, especially with respect to *Pelomonas*. Collectively, future research should seek to establish the precise route by which *Pelomonas* synthesizes salicylaldehyde to affect IHP infections and whether related mechanisms exist among the microbiota of other vertebrate hosts.

Our multi-omic analysis reveals substantial alterations in NAEs and NAE precursors between parasitic infected and uninfected hosts, with a potential route of control by the gut microbiome. Given that NAEs play in a broad range of physiological functions such as immune regulation as well as energy metabolism and feeding behavior^84–86,108,109^, IHPs are capable of modulating NAEs to enhance infection^90^, and their alteration is associated with changes in the gut microbiome^87^, these compounds represent a potentially rich area to understand the intersection of parasite infection with the gut microbiome and host immune regulation. Our results demonstrate that alteration of the gut microbiome by antibiotic exposure appears to drive changes in NAE abundance that are sustained in the profile of several NAE compounds several weeks after initial antibiotic exposure. The abundance of these compounds is also linked to IHP infection and is linearly related to IHP infection burden. The NAE axis represents one route by which the intestinal microbiome drives aspects of host physiology. The functions which are regulated by NAE signaling such as feeding behavior and inflammation response are likely relevant in explaining some aspects of parasite infection, especially given that parasites also possess enzymes involved in regulating NAEs and their associated physiological functions. The synthesis of these findings is complex, but elucidating the principal effects of NAE changes on host physiology and parasite infection, in addition to identifying taxa whose abundance is related to NAE changes, may reveal how intestinal microbiota participate in mediating host response to IHPs and could reveal drug or probiotic targets that aid in control of helminth infection.

While we highlight several compounds, such as salicylaldehyde and NAEs, which help explain the relationship between the gut microbiome and parasite infection, there exists a rich unexplored repository of metabolites in this data that may display similar anthelmintic activity. To better understand the metabolomic landscape relevant to IHP infection, we modeled the relationship between parasite burden and metabolite abundance and elucidated a collection of compounds that may be harnessed and further investigated in efforts to control parasitic colonization and success. For instance, trunkamide A, a compound that is of known bacterial origin^110^, and which has been examined for its antibiotic and antitumor activity^111^, shows up as being inversely associated with parasite burden in our data. Given the established ability of this compound to be produced by bacteria, understanding its distribution among aquatic and gastrointestinally associated bacteria may offer an enticing route to the discovery of novel probiotic microbes which could be supplemented and stimulated to produce trunkamide A under specific parasite-related conditions. Additionally, several compounds which have been explored for anthelmintic activity, including baliospermin^112^, and genistin^113^, also show up in our results as being significantly linked to parasite burden. Some of these metabolites may be of bacterial origin, or modified by members of the gut microbiota, such as genistin^114^, in a manner that enhances their anthelmintic activity. Applying mediation methods to understand the possible relationships between these compounds and microbes involved in parasite infection may help to establish microbe-dependent routes for the control of parasite infection. Collectively, these findings represent an additional set of metabolomic compounds that may be explored by mining the gut microbiome for potential solutions to the urgent challenge posed by increasing anthelmintic drug resistance.

In summary, this work unravels interactions within the zebrafish gut ecosystem, yielding a deeper understanding of the dynamic relationships among microbiota, metabolites, and parasitic infections. These results extend the application of mediation inference methodologies to reveal specific bacterial metabolites that may serve as key mediators of host-parasite interactions, and identifies novel anthelmintic drug leads. Notably, salicylaldehyde emerges as a compelling anthelmintic compound, and we demonstrate its ability to inhibit parasitic egg maturation in zebrafish. This work also establishes the involvement of other metabolites, like N-acylethanolamines, in host-parasite-microbiome dynamics emphasizing the need for further research to elucidate their influence. Collectively, our findings support the hypothesis that gut microbiota play a role in parasite infection, and understanding the chemical means by which microbiota are involved in helminth colonization may yield tools for infection control.

## Methods

### Zebrafish Husbandry, Parasite Exposure, and Parasite scoring

All zebrafish research was conducted under the approval of IACUC protocol 2022-0280. Tropical 170 day old 5D zebrafish were obtained from the Sinnhuber Aquatic Research Laboratory (Corvallis, OR). The fish were housed in a flow-through vivarium on a 14hr:10hr light:dark cycle, and fed Gemma Micro 300 once a day, except on the weekends. Water temperature was recorded daily and ranged from 23-28 C. There was also weekly testing of ammonia (0-0.25ppm), pH (7.6), hardness (0-25 ppm), and conductivity (90-110 uS/cm) to ensure high water quality. Prior to initial antibiotic exposure or parasite exposure, fish were allowed to equilibrate in their tanks for 2 weeks. Each of the 100 fish used in this protocol was randomly assigned to one of four unique exposure groups, no parasite and no antibiotic exposure, no parasite and antibiotic exposure, parasite exposure and no antibiotics, and both parasite and antibiotic exposure. After this period of equilibration, 50 out of the 100 fish were exposed to a combination of antibiotics, 10mg/L colistin and 10mg/L vancomycin. Of the 100 fish used in the experiment, 50 were exposed to *P. tomentosa* eggs at a dose of 88 eggs per fish.

### Zebrafish Gut Metabolite Mass Spectrometry

Fecal pellets collected from individual zebrafish were promptly lyophilized to minimize leaching of metabolomic content into water and then split approximately equally for use in metabolomic and 16S rRNA library preparation and sequencing analyses. Due to the small sample size and high water content of the fecal pellets, precise weighing was not feasible. Therefore, total ion abundance normalization was implemented to account for variation in sample input. Ivermectin and two isotopically labeled amino acids were incorporated into the extraction mix to monitor injection accuracy and platform performance throughout the extended batch run time. Zebrafish fecal samples were prepared for untargeted metabolomic analysis using a modified extraction protocol. An extraction solvent consisting of equal parts 100% ethanol (Sigma 1.11727.1000) and methanol (Fisher A456-1) was prepared and chilled overnight at −20°C. Three 1.4mm zirconium oxide beads (VWR 10144-554) were added to each 2ml screw-top bead beating tube (Fisher 02-682-558). An extraction mix was then created by diluting isotope standards (Cambridge Isotope Laboratories MSK-A2.1.2; 1:20 dilution) and 100µM Ivermectin standard (Sigma PHR1380) 1:10 in the chilled ethanol/methanol solution. Twenty-five microliters of this extraction mix were added to each tube containing a fecal sample. The samples were then homogenized using a Precelly’s 24 bead beater (program: 2 x 5400 rpm for 45s, 5s wait interval), centrifuged at 16,000 x g for 10 minutes at 4°C, and the resulting supernatant (15-25µL) was transferred to glass vials (Microsolv 9532S-3CP-RS). In cases of large sample volumes, an additional centrifugation step was performed.

Extracts were submitted for analysis using untargeted LC-HRMS/MS-based metabolomics. An AB Sciex TripleTOF 5600 mass spectrometer coupled to a Shimadzu Nexera UHPLC system was used as described previously^115^. Metabolite extracts were separated using an Inertsil Phenyl-3 column (2.1 x 140 mm, 100 Angstrom, 5 µm; GL Sciences, Torrance, CA, USA). Column was held at 50°C. We used a 50-minute binary gradient system consisting of: Solvent A, water (LC-MS grade) with 0.1% v/v formic acid and solvent B, methanol (LC-MS grade) with 0.1% v/v formic acid. Metabolites were eluted using the following gradient program: 1 min, 5% B; 11 min, 30% B; 23 min, 100% B; 35 min, 100% B; 37 min, 5% B; 37 min, 5% B and 50 min, 5%B. Flow rate was 0.5 mL/min. Injection volume was 10 µL. Positive IonSpray voltage was set to 5200 V, negative IonSpray voltage was set to 4200 V. Source temperature was 350°C. The q-TOF mass spectrometer was operated in the data-dependent mode using the following settings: period cycle time 950 ms; accumulation time 100 ms; m/z scan range 50– 1200Da; and collision energy 40 V. Mass calibration of the TOF analyzer was performed automatically after every fifth LC run.

LC-HRMS/MS data processing was performed with Progenesis QI software V2.0 (NonLinear Dynamics, United Kingdom) and ABSciex Masterview (ABSciex, USA) entailing peak picking, retention time correction, peak alignment, and metabolite identifications/annotations. Metabolite annotations was facilitated by Progenesis QI and Masterview using an in-house spectral library base on the IROA Mass Spectrometry Metabolite Library of Standards (MSMLS) containing retention times, exact mass and MS/MS information of >650 metabolite standards (IROA Technologies, Bolton, MA, USA). This workflow allows obtaining high confidence annotations (L1). In addition, tentative metabolite annotations were obtained by searching the METLIN MS/MS library in Progenesis QI.

### Microbial 16S rRNA Library Preparation and Sequencing

Fecal samples were collected from individual adult fish at Days 0, 3, and 32. Fish mortality and parasite exposure precluded the collection of fecal pellets from every fish at all time points. For samples and time points at which fecal samples could be collected, DNA was isolated using the Qiagen DNeasy PowerSoil kit, in accordance with the manufacturer’s directions. After a 10-minute incubation at 65°C, samples were subjected to bead beating, using 0.7mm garnet beads, for 20 minutes using the Vortex Genie 2 (Fisher, Hampton, NH, USA). PCR was performed in triplicate using 1 microliter of purified DNA from the lysis solution to amplify the V4 region of the 16S rRNA gene, using the 806r and 515f primer set^116^. Amplified DNA collections were quantified using a Qubit HS Kit (Carlsbad, CA, USA). An equal quantity of DNA was selected from each of the 252 samples, for a total DNA mass of 200ng, and the pooled collection of DNA was cleaned using the QIAGEN QIAquick PCR purification kit then diluted to a final concentration of 10nM. The final pooled DNA collection was sequenced by the Center for Quantitative Life Sciences at Oregon State University, using an Illumina MiSeq instrument with 250-bp paired-end reads.

### Zebrafish Gut Microbiome Community Diversity Analyses

Read quality filtering was performed using DADA2^117^ with R 4.1.2. Alpha and beta-diversity analyses were performed using a relative abundance-normalized sequence count table. Generalized linear mixed effects models were used to model species richness and Shannon diversity as a function of longitudinal timepoint, fish id, antibiotic exposure, parasite exposure, and an interaction of antibiotic and parasite exposure. Bray-curtis dissimilarity and subsequent NMDS ordination was performed using vegan^118^. To clarify how antibiotic administration, parasite exposure, and other host factors relate to gut community composition we used permutational multivariate analysis of variance (PERMANOVA, adonis2, vegan) with 10,000 permutations.

### Mediation analysis of gut bacteria and metabolites

Regression-based mediation analysis was used to investigate the hypothesis that the relationship between fecal metabolites to nematode parasite burden is mediated by the abundance of gut microbiota. Briefly, the standard approach for these techniques employs regression modeling to analyze the association between two variables, then models the possible effect of a mediating variable on the relationship between the variables in the initial model. We constrained the initial feature space by selecting metabolites for which annotation and research-characterized biological identity was available. Sparsity of gut bacterial data challenges statistical investigation of correlation, so only taxa whose relative abundance was greater than 0 in more than 30% of samples were used. Microbiome data was normalized using relative abundance, and the log of metabolite MS abundance was log transformed adding 1 to initial values which were equal to 0. Initial feature selection yielded 40 prominent bacterial taxa, and 27 metabolite compounds which were significantly related to worm burden (NB-GLM, FDR < 0.1). We utilized the nptest package^119^ to test for partial correlation for pairwise relationships between metabolites and bacterial taxa. We applied partial correlation for each microbe-metabolite pairing, and used all remaining taxa as conditioning variables to isolate direct microbe-metabolite links. Pairings with an FDR-adjusted relationship <0.3 were considered. Concurrently, ASV mediation of metabolite-parasite burden was tested using the mediation package^120^ in R, where metabolites were coded to represent high and low metabolite abundances, with ASVs as mediators. Average causal mediation effect (ACME), average direct effect (ADE), and proportion of direct effect mediated were calculated for each model of microbe-metabolite pairing. Family-wise error rates were controlled by applying Benjamini-Hochberg FDR correction (FDR<0.3) to partial correlation and ACME p-values for tested mediating relationships. Visualizations of putative mediating interactions were visualized using ggplot2^121^. Code to recreate the mediation analysis and visualizations is available here (https://github.com/CodingUrsus/Zebrafish_Microbiome_and_Parasites).

### Salicylaldehyde Toxicity Assay

A toxicity assessment identified the highest SA dose which did not result in significant mortality. Adult zebrafish were aqueously exposed to SA in two 48 hr periods spaced 5 days apart. The fish were monitored for any adverse health effects including mortality, and fish were evaluated to ensure they were alive and capable of active movement. Toxicity endpoints were evaluated from the initial exposure to 7 days after the last exposure. The chemicals used in this study were salicylaldehyde (SA, cas: 90-02-8) and dimethyl sulfoxide (DMSO, cas: 67-68-5). Salicylaldehyde was obtained from Tokyo Chemical Industries (lot: J052M-EQ) and DMSO was purchased from VWR (lot: 22H2456964). Dilutions were made with 100% DMSO and stored in closed vials in a desiccator at room temperature.

Fish were aqueously exposed to 0,1, 2, 3, 5, and 10 mg/L of SA with 0.01% DMSO. The 0-2 mg/L groups had two replicate tanks containing 16 males and 16 females. Furthermore, the 3-10 mg/L groups had 1 replicate tank containing 6 males and 6 females. The 3-10 mg/L exposure groups were intended as positive controls, to ensure that null effects were not a result of no chemical exposure.

Exposure groups 0-2 mg/L SAL occurred in 9L tanks first filled with 3L of system water followed by 1mL of a concentrated SA stock solution. Following the addition of SA, the remaining 3L of fish water was added to the tank to mix the chemical. The 3-10 mg/L exposures were conducted in 2.8L tanks and followed the same chemical additional pattern. However, only a total of 2L of water was added to the tank. Solutions were refreshed every 24 hrs using the same method previously described. Lastly, each exposure tank was aerated with an air stone and the fish were not fed, to preserve water quality.

### Parasite Salicylaldehyde Exposure Assays

#### a. *in vivo* Salicylaldehyde Exposure

Based on our previous transmission studies^49,102^, we experimentally infected 100 5D line fish by placing them in a large tank from which ∼30 infected fish were removed the day before. The recipient fish were exposed in the infection tank for 5 days. *P. tomentosa* infection was promoted within the tank by reducing the waterflow, not removing the detritus, and keeping fish carcasses in the tank. Over a 5-day period, 1L of water from another infected tank was added to the exposure tank to further enhance infection. After 5 days of exposure, the remaining fish (∼75) were randomly divided into four 9L tanks, two control tanks and two salicylaldehyde exposure tanks.

Fish were then exposed to salicylaldehyde in the same manner as the toxicity trials in their aquaria with the water turned off at 14-15 and 21-22 days post initial exposure (dpe). Using the results of the toxicity trial, fish were dosed with either 2mg/L SA with 0.01% DMSO or 0.01% DMSO (controls). Fish were monitored daily, and fresh dead fish were examined for the presence of the worm. At 24 dpe, the infection status of the fish was evaluated using wet mounts. The fish were first euthanized using a hypodermic shock. Following euthanasia, the gut was dissected out of the fish and placed on a glass slide with a 60 × 24 mm coverslip with about 200 µL of water. The gut was compressed with a cover slip and viewed on a compound microscope at 50 and 100x magnification. While viewing the guts the number of immature worms, mature female eggs were counted. Each slide was read by two different readers within about 20 min., and the average worm counts were used for future analyses.

#### b. Egg Larvation Assay

We tested the ability of SA to inhibit egg larvation through two separate trials in line with previously used protocols^102^. Briefly, *P. tomentosa* eggs were collected by placing fish in a 10L static tank for 2 days. After two days, the water was filtered through a custom 3D printed filter apparatus fitted with 105, 40, and 25 µm nylon screens. The material retained on the 25 µm screen was collected in 15 mL conical tubes and centrifuged at ∼3,000 x g for 5 minutes. The supernatant was decanted until 1ml of water remained.

In a first trial, collected eggs were exposed to 0, 2 or 15 mg/L of salicylaldehyde with 0.01% DMSO in 15 mL conical tubes. Eggs were transferred into each exposure group by pipetting 200 µL of the egg pellet formed as described above into each conical tube. The solution was homogenized and transferred to a 30°C room for incubation. *P. tomentosa* egg larvation was quantified after 5 days of SA exposure (∼10 days old). A second trial was conducted where eggs were placed in a container on a shaker to aid in distribution of DMSO control and SA throughout the egg-hatching solution. Eggs were exposed to 0, or 2 mg/L of SA with 0.01% DMSO in 15 mL conical tubes, with the shaker on speed setting 2.5 (Hoefer, San Francisco CA) throughout the exposure. Larvation of eggs was quantified 3 days after SA exposure (∼6 days old). In both trials, after exposure eggs were collected by centrifuging the tubes at 5,000 rpm for 5 minutes. The eggs were subsampled by taking 20µL of water from the bottom of the tube and placing them on a glass slide and covered with a glass 24×24 mm coverslip. The number of larvated and unlarvated/dead eggs were counted using a compound microscope at 50, 100 or 400x magnification.

## Acknowledgements

We would like to express our gratitude to Kristin Kasschau and Alexandra Alexiev for their valuable insights on microbial growth and metabolism requirements.

This project was supported by NIH NIAID R21 to TJS (R21AI135641), NIH NIEHS R01 to TJS (R01ES030226), NSF grant (2025457) to TJS, NIH grant (S10RR027878) to JFS, and a Tartar fellowship to AJH.

## Data Availability

The nucleotide data underlying the findings of this study are available in the NCBI Sequence Read Archive (SRA) under BioProject ID PRJNA1132310, and annotated metabolomic data from positive and negative ion modes are available here (https://github.com/CodingUrsus/Zebrafish_Microbiome_and_Parasites/).

---

Supplementary Figure 1. Distribution of mature *Pseucapillaria tomentosa* worms quantified during dissection of intestinal tissue 29 days after initial helminth egg exposure.

Supplementary Figure 2. The ten most important features based on the increase in node purity for regression of helminth worm burden measured 29dpe on microbiota relative abundances at 0dpe, just prior to parasite egg exposure.

Supplementary Table 1. Coefficients table of metabolites which are linked to IHP burden (FDR<0.1) as measured 29dpe.

Supplementary Table 2. Coefficients table of PERMANOVA results testing the relationship between NAE abundance and microbiome composition prior, as well as the interaction of NAE abundance and prior antibiotic exposure at 0dpe and 29dpe.

