## Supplemental Tables for "Gut microbiota metabolically mediate intestinal helminth infection in Zebrafish"

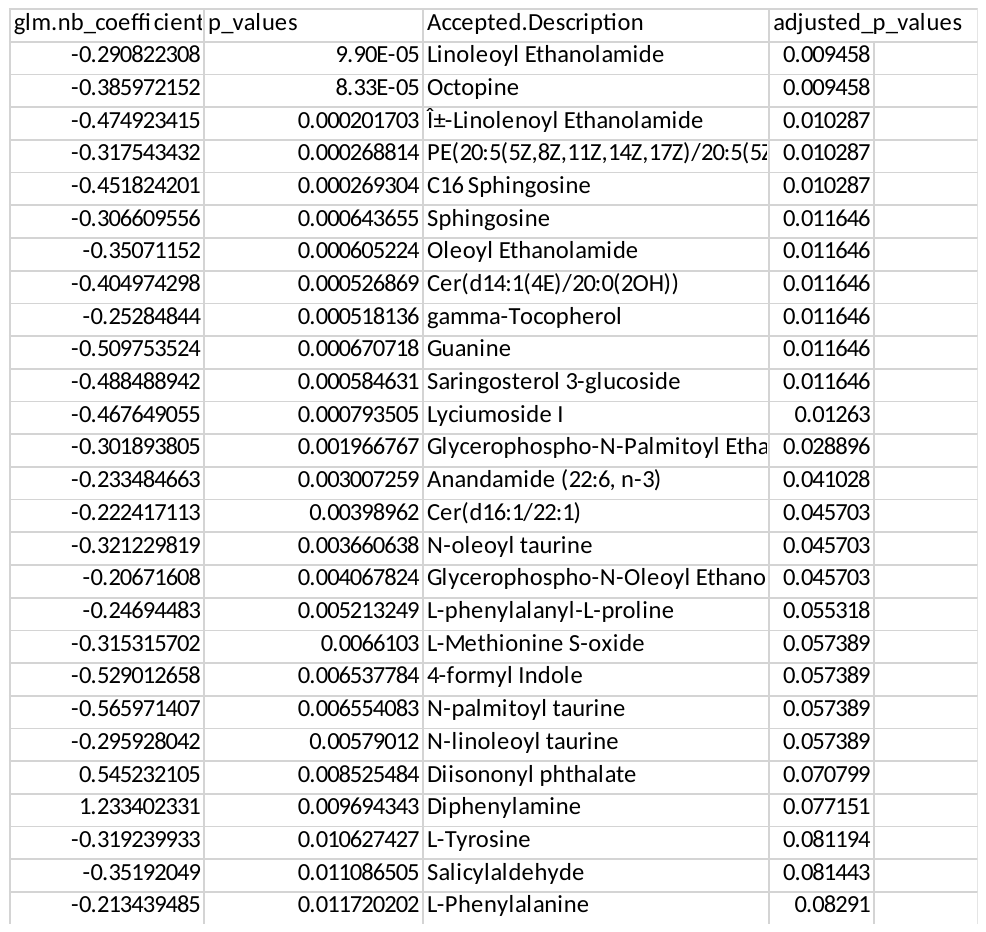

Supplementary Table 1. Coefficients table of metabolites which are linked to IHP burden (FDR<0.1) as measured 29dpe.

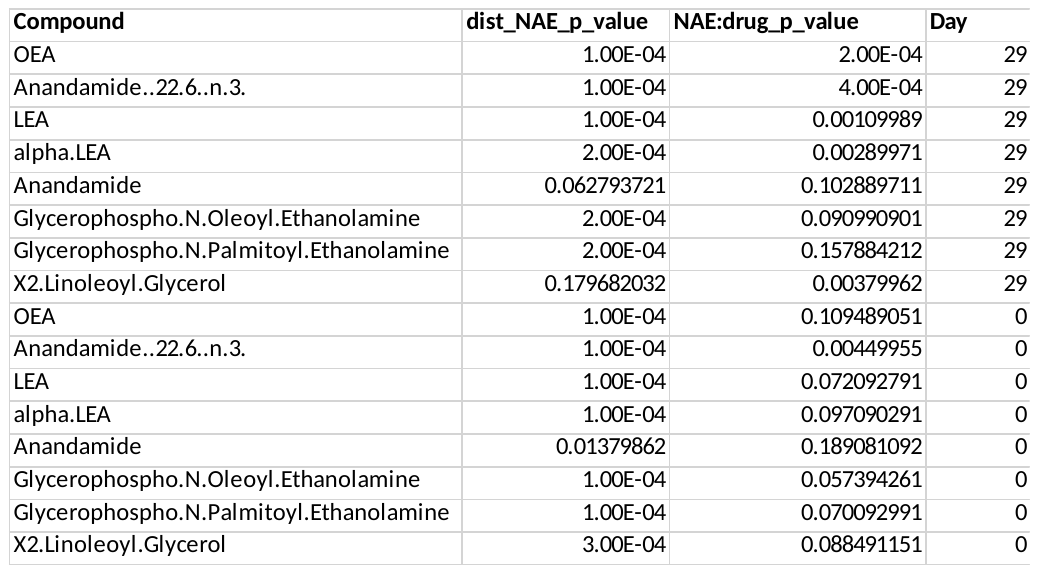

Supplementary Table 2. Coefficients table of PERMANOVA results testing the relationship between NAE abundance and microbiome composition prior, as well as the interaction of NAE abundance and prior antibiotic exposure at 0dpe and 29dpe.
